# Raman mapping glucose metabolites during human mesenchymal stem cell adipogenesis

**DOI:** 10.1101/2020.04.19.048827

**Authors:** Gomathy S. Subramanian, Con Stylianou, In Yee Phang, Simon Cool, Victor Nurcombe, Martin J. Lear, Sergey Gorelik, David G. Fernig, Jonathan Hobley

## Abstract

Raman mapping was used to determine the lipid distribution inside human mesenchymal stem cells during induced adipogenesis by monitoring C-H stretching bands of the fats inside the lipid droplets. By incorporating deuterated glucose into the cell culture medium during induction it was possible to distinguish whether or not downstream metabolites, either in lipid droplets or in the cytoplasm, had been formed before or after the adipogenic cascade, because C-D stretching bands are 1/√2 shifted compared to the C-H bands. Thus, metabolites formed after the initiation of the process displayed both C-H and C-D stretching bands and so were forming during induced adipogenesis rather than prior to it. With the ability to distinguish small putative lipid drops formed by the induction of adipogenesis from those pre-formed in the cell, it was possible to analyze spectral changes occurring in the droplets at the earliest stages of adipogenesis. There were two key findings. Firstly, Raman spectra of lipid droplets evolved over time, suggesting that their composition at the early stages was not the same as at the later stages. Secondly, it was apparent that the proportion of unsaturated fats in droplets was higher at early stages than it was at later stages, suggesting that unsaturated fats arrive in the droplets faster than saturated ones.

## Introduction

Adipocytes are cells that convert and store excess energy in the body as lipid droplets. They are also involved in secreting a host of regulatory factors that are important for the maintenance of normal metabolism (1). Understanding the processes behind glucose metabolism, adipogenesis and lipolysis is thus important for the understanding of conditions such as obesity and diabetes. The formation and growth of lipid droplets have been studied using techniques such as mass spectrometry, fluorescence labeling and protein extraction assays, both *in vitro* and *in vivo* (2–6). Although these methods have allowed insight into the process of adipogenesis, two key questions remain unanswered, how the droplets actually grow, and where they initially form (5).

Raman spectroscopy is a technique that can be non-invasive and that requires minimal sample preparation, but which enables a characterization of both the morphology and the biochemical content of live cells (7–11). Additionally, more recently coherent anti-Stokes Raman scattering (CARS) has become popular for cell imaging (12–17), albeit it presents great technological challenges. Although in its infancy in this field (12,14–16) we believe that Raman mapping can contribute greatly by providing an *in situ,* label-free method for observing biochemical reactions in living cells.

In spontaneous Raman spectroscopy, inelastic scattering results in scattered light that is frequency shifted from that of the incident laser light. This frequency shift depends on the vibrational modes available in the chemicals that the light is scattered from. Hence these shifts can be used to quantify the vibrational energies of the different chemical bonds in the medium being studied. Each molecule or group of molecules, therefore, has its own “fingerprint” Raman spectrum that is based on its chemical bonds, hydrogen bonds and solution conformations. Molecules in cells such as proteins, DNA, carbohydrates and lipids have all been identified using this technique (8–11,18,19). Additionally, coupling a Raman spectrometer with a confocal microscope allows imaging of vibrational spectra of live cells. With such a system, mapping the spatial differences in biochemical composition within the cell becomes possible.

Here we describe the use of Raman microscopy to study adipogenesis, and the use of deuterium labeling to specifically track the D-glucose-d7 metabolites formed during the differentiation of this lineage from human mesenchymal stem cells (hMSCs). Adipocytes provide an almost ideal system to study using this method because, during their maturation, lipid droplets form that contain high concentrations of fats, which overcomes a common limitation of Rama microscopy: the signal from the many dilute molecules in complex cytosolic mixtures in the cells is weak (11). This means that the lipid droplets can be clearly located by their very intense C-H bands from ~2800 cm^−1^ - 3000 cm^−1^. In this case, a bulk medium in droplets (the lipid droplet) is observed, which is itself within another bulk medium, as the continuous phase (the cytoplasm). Furthermore if d7-deuterium labeled D-glucose is added to the culture medium for the cells, it is possible to further specify the regions of the lipid’s molecular skeleton that can be interrogated. This is because deuterated glucose and its derivatives have a C-D band that is shifted from the naturally abundant C-H band by a factor of 1/√2, thus enabling the spectral changes to be seen as the D-glucose-d7 is converted to the triacylglyceride (TAG) backbone. It is important to note that D-labeling of glucose is particularly non-intrusive, making it possible to track the glucose and its metabolites without further labeling.

## EXPERIMENTAL PROCEDURES

### Cell culture

hMSCs (Lonza) were cultured in Dulbecco’s modified Eagle’s medium (DMEM 1mM sodium pyruvate (GIBCO (Invitrogen)) and 4mM L-glutamine (GIBCO (Invitrogen)) added just before culture) with 1 mg/ml glucose supplemented with 10% (v/v) fetal bovine serum (FBS, HyClone) and penicillin-streptomycin (100 IU/ml each, GIBCO (Invitrogen)). They were seeded (5000 cells/cm^2^) onto sterile glass coverslips (10mm diameter Nunc Thermanox glass Coverslips from Thermo Fisher Scientific) and grown to confluence. An adipogenic medium consisting of DMEM (low glucose, with 1 mM sodium pyruvate (GIBCO (Invitrogen)) with 4.5mg/ml glucose (either D-glucose or D-Glucose-1,2,3,4,5,6,6-d7, 97 atom % D, Sigma-Aldrich St Louis MO USA) 10% (v/v) FBS, penicillin-streptomycin (100 IU/ml each), methylisobutylxanthine (IBMX, 115μg/ml), indomethazine (20 μM) 1μM dexamethasone (all from Sigma Aldrich) and insulin (1.7 μM) was used to promote the differentiation of the hMSCs to the adipocyte lineage. Cells on glass coverslips were immersed in 1x PBS buffer for imaging using the Raman microscope.

Batches of cells were also similarly cultured for analysis of the D-glucose and total fat content in the growth medium during adipogenesis. This was done to evaluate the concentration of the main source materials for adipogenesis over the timescale of the process. To do this, a commercially available glucose meter was used to determine the glucose content (Xceed Blood Glucose Monitor, Abbott Diabetes Care Inc., USA and MediSense Optium blood glucose electrodes and strips to load sample, Abbott Diabetes Care).

The fats were determined by their weight after extraction. Medium (700 μl) was put into an Axygen centrifuge tube and 770 μl chloroform: methanol (1:2) added. The suspension was vortexed for 1 min and then incubated on ice on a shaker for 30 min. It was then centrifuged (14000 rpm Eppendorf 5417R) for 2 min at 4°C. The bottom organic layer was then transferred into a fresh tube. The sample was dried partially under vacuum (2000 rpm Scanvac Minivac Alpha vacuum condenser, 20 min). To the remaining aqueous phase, 105 μl of chloroform was added. This was then vortexed hard for 2 min, centrifuged (14000 rpm Eppendorf 5417R) for 2 min and then transferred to the partially dried tube. The sample was then dried to completion under vacuum (2000 rpm Scanvac MiniVac Alpha vacuum condenser, 50 min). The lipids were stored in a −80°C freezer until the analysis of their mass.

### Raman mapping

A confocal Raman microscope system (Witec, Ulm, Germany) was used to obtain Raman spectra and spectral maps. This system has a 532 nm laser coupled into a single-mode fiber attached to the microscope. A 60x water immersion objective (1.0 NA, Nikon) focused the laser beam onto the sample. The laser spot was scanned across the sample using a piezoelectric motorized stage, collecting a spectrum from every pixel in a user-specified region. The Raman signal from the sample was captured through the same objective and sent through an edge filter to suppress Rayleigh scattering, before being coupled into a multimode fiber. This enabled Raman scattered light from only the focal plane to be collected and analyzed by a reflective grating spectrometer with a back-illuminated CCD camera. Raman maps were created for a number of different cells on each day of measurement from before and after induced adipogenesis in order to make sure that we analyzed a representative selection of cells over time. Different cells were analyzed on different days.

Witec software was used to analyze the spectral maps. A Raman image for specific vibrational bands was created by selecting an interval of wavenumbers and displaying a map of the integral intensity of the band for every pixel. The C-H band images display the integrated intensity in the range of 2820 to 3020cm^−1^, and C-D band images were integrated in the range of 2080 to 2280 cm^−1^. Cluster analysis of spectral maps was performed using Witec software.

## RESULTS AND DISCUSSION

### Human MSCs

Raman images of live hMSCs were obtained for comparison with the images of hMSCs differentiated towards the adipocyte phenotype into which they developed at later stages after suitable induction. A typical optical and Raman image, which maps the spectral region from 2830cm^−1^ to 3007cm^−1^ for an hMSC, is shown in **Fig. 1**. In this map we can see clear details of the cell, which present as intensity variations that conform to the cell’s morphology. Spectra of hMSCs were averaged over several regions of the cell, using cluster analysis, and subjected to a detailed comparison of the various chemical signatures. Assigning a cellular region based on a Raman image is particularly complicated, because without fluorescent labeling with commercially available markers (which mask Raman signals), identifying cell components reliably is difficult. It is, therefore, necessary to first review spectral assignments from previous studies to be able to compare them with our data; **Table 1** summarizes this information.

**Table 1.** Peak assignments for Raman bands measured in hMSCs.

| Band<br>(present data) | Band<br>(reference data) | Assignments | Comment | Reference<br>(respective) |
| --- | --- | --- | --- | --- |
| Low Frequency Region |  |  |  |  |
| 720 | 719; 719 | Choline (H<sub>3</sub>C)N<sup>+</sup> | W | 8, 18 |
| 790 | 785; 788; 788; 788 | Nucleic acids cytosine, thymine, uracil; C ring breathing, thymine / nucleotide backbone PO<sub>2</sub> sym str | M | 7, 8, 18, 19 |
| 945(S) | 937; 936; 938 | Protein α-helix and C-C skeletal modes | | 8, 18, 19 |
| 1010 | 1003; 1005; 1005; 1004 | Protein C-C, Phe | S, Sharp | 7, 8, 18, 19 |
| 1070 | 1060-1095; 1065 | Chain C-C | W | 8, 18 |
| 1096 / 1099 | 1093; 1060-1095; 1094-1100; 1094 | Nucleotide backbone (PO<sub>2</sub>) | M | 7, 8, 18, 19 |
| 1130 | 1126; 1128; 1128; 1126 | C-C stretch; protein C-N str | M | 7, 8, 18, 19 |
| 1256 (S) | 1220-1284; 1255; 1254 | Protein amide III; adenosine | W | 8, 18, 19 |
| 1277 | 1263; 1220-1284 | T,A Nucleic acids; C-H bend, Amide III Proteins; =CH bend Lipids |  | 7, 8 |
| 1302 | 1302, 1301; 1300; 1304 | CH<sub>2</sub> twist lipids; Protein amide III; adenosine |  | 7, 8, 18, 19 |
| 1331 | 1337; 1342; 1340; 1339 | Proteins C-H def; Adenosine; Guanine |  | 7, 8, 18, 19 |
| 1348 |  |  |  |  |
| 1446 / 1455 | 1447; 1449; 1451 | CH<sub>2</sub> def; C-H def |  | 7, 18, 19 |
| 1580 | 1575; 1577; 1578; 1578 | Guanine, Adenosine |  | 7, 8, 18, 19 |
| 1616 |  |  |  |  |
| 1664 | 1655-1680; 1660 | C=C str. Lipids, Protein amide I |  | 7, 8, 18, 19 |
| High Frequency region |  |  |  |  |
| 2729 |  |  |  |  |
| 2856 | 2854; 2854 | CH<sub>2</sub> Str Lipid |  | 18, 19 |
| 2890 | 2886; 2886 | CH<sub>2</sub> Str Lipid |  | 18, 19 |
| 2940 | 2936; 2936 | CH<sub>3</sub> Str |  | 18, 19 |

**FIGURE 1.**
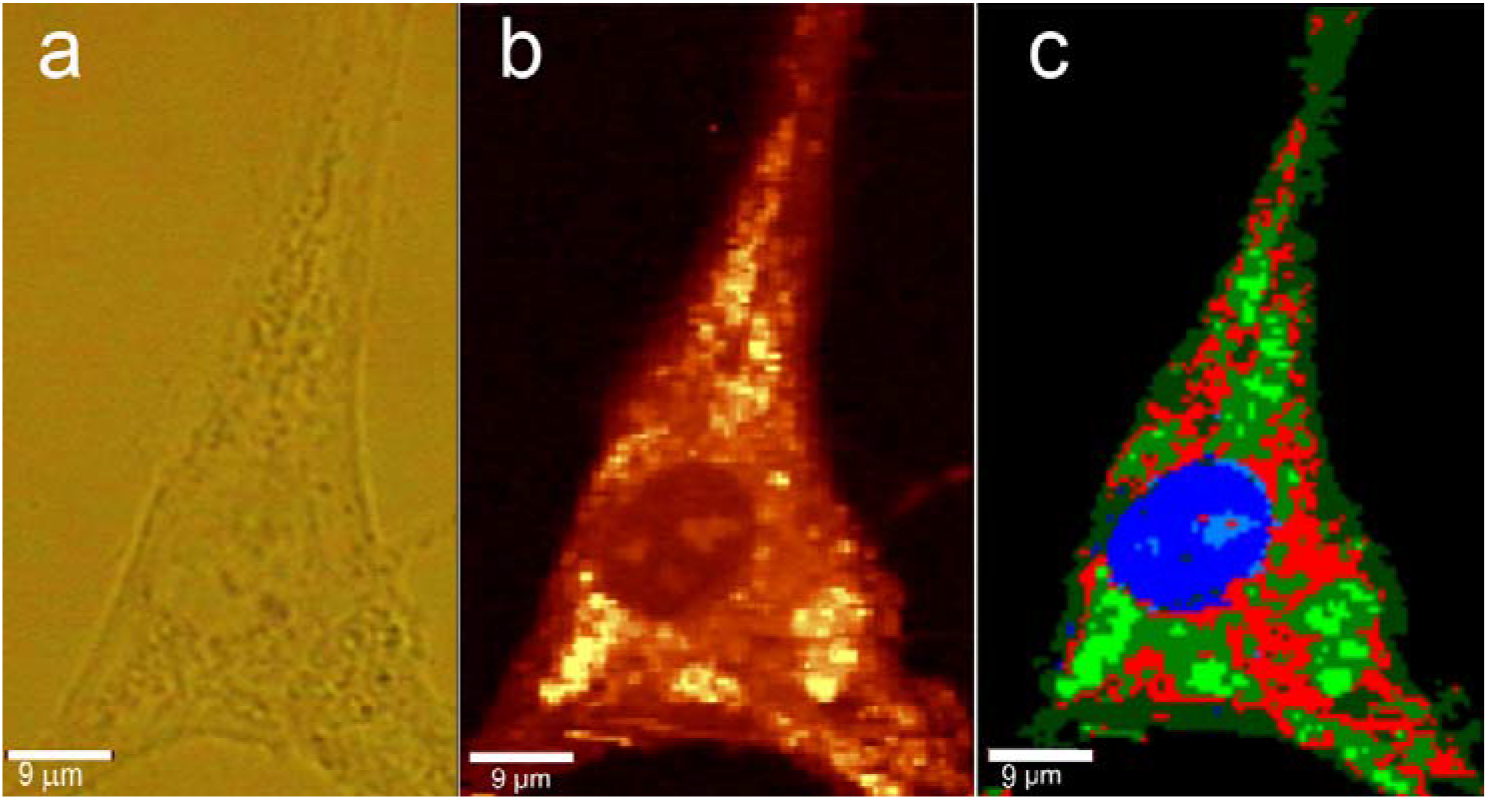
**a)** Optical image of an hMSC. **b)** The corresponding C-H stretch Raman map. **c)** Cluster analysis of the same hMSC. Blue and lighter blue corresponds to the nucleus and chromatin complex, respectively, having a spectrum corresponding to that of DNA and chromatin, shown in Figure 2(a-b). Green corresponds to brighter regions outside the nucleus and has spectra corresponding to spectrum 1 in Figure 2(c-d)). Red corresponds to the less bright regions outside the nucleus and corresponds to spectrum 2 in Figure 2(c-d). Dark green corresponds to the darkest regions outside the nucleus and has spectrum similar to 3 in Figure 2(c-d).

The area of the hMSC Raman maps that could be most easily identified was the nucleus (**Fig. 1)**. Within the nucleus, brighter areas could be observed that may be assigned to the chromatin complex. Spectra of the brighter and darker areas of the nucleus were very similar in terms of the peaks that could be seen (**Fig. 2**). However the brighter regions had a slightly greater contribution from the DNA bands at 790 cm^−1^, 1099 cm^−1^ and 1580 cm^−1^ (8,18,19).

**FIGURE 2.**
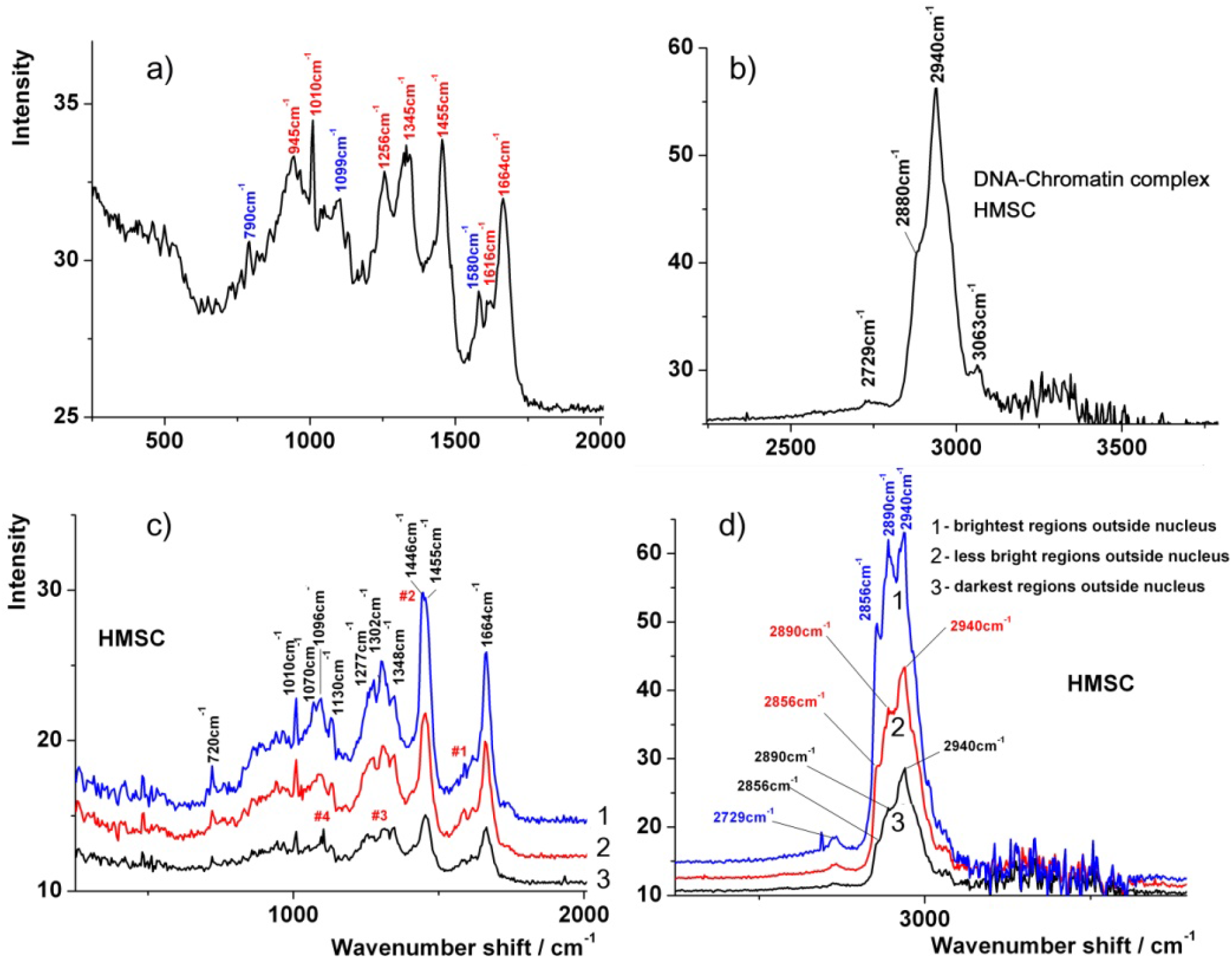
Raman spectra of hMSCs. Top:**a) and b)** The spectrum of the nuclear DNA-chromatin complex was averaged over 4 different cells. Bottom: **c) and d)** spectra derived from the cluster analysis in Fig. 1.

From previous studies, the peaks at 790 cm^−1^ can be assigned to the phosphate backbone of DNA or to cytosine, thymine and uracil. Peaks at 1099 cm^−1^ can be assigned to the nucleotide backbone and 1580 cm^−1^ can be assigned to guanine and adenosine. Peaks at 1256 cm^−1^ and 1331 cm^−1^ can be also assigned to adenosine (see Table 1), as they are seen in the nuclear spectrum, but not in the spectra obtained outside the nucleus (18,19).

All other peaks in the low frequency region can be assigned to the large amounts of protein present in the nucleus. Few lipids were anticipated in the hMSC nucleus, and this was confirmed by the spectrum in the C-H stretching region, which did not reveal any of the characteristic lipid peaks at 2856 cm^−1^ and 2890 cm^−1^ (10,18). Thus the C-H band obtained from the nucleus is most likely to arise from proteins and nucleic acids.

Outside the nucleus it is much harder to assign different regions to specific classes of molecule. However, some regions were clearly brighter than others. Cluster analysis could be used to identify different spectral regions, using spectra extracted from these regions. From this it could be seen that the brightest regions of the image corresponded to regions with spectra showing characteristics of higher lipid content. Despite the fact that several C-H bands overlap in the region 2800 cm^−1^ to 3000 cm^−1^, it is possible to identify lipids by the increase in the intensity of CH_2_ stretching bands at 2856 cm^−1^ and 2890 cm^−1^ (**Fig. 2**) (10,18). These bands were clearly more prominent in the brighter regions outside the nucleus. Additional changes in the spectra in lower frequency regions could also be accounted for by contributions from lipid bands, as can be seen by comparison to the spectrum of a lipid droplet from a 30-day old adipocyte shown in **Fig. 3.** The only other difference between spectral regions that was notable was the presence of the 1580 cm^−1^ peak for adenosine and guanine in the spectra obtained from both the less bright and the darkest regions outside the nucleus, which may arise from RNA and/or the bases themselves.

**FIGURE 3.**
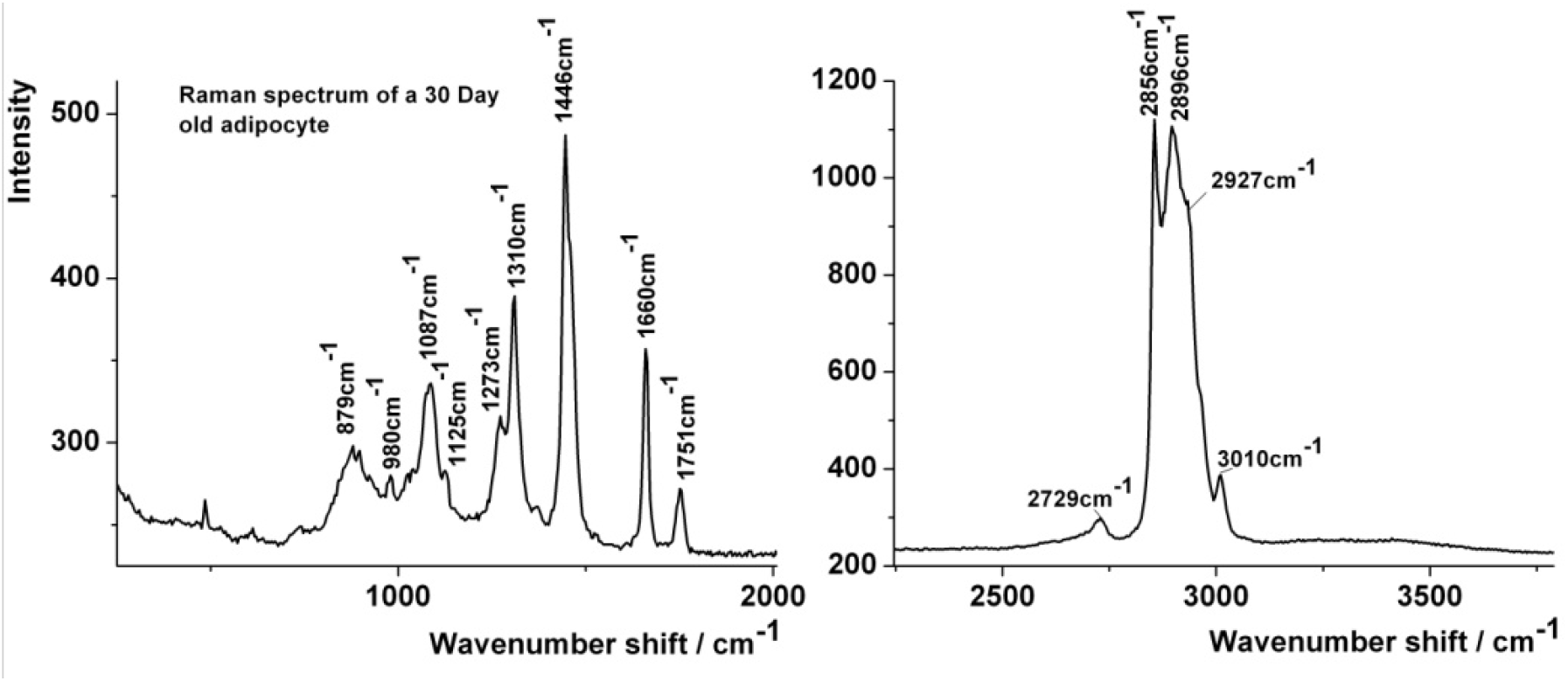
Spectrum taken from the centre of a large (>10μm) lipid droplet in an adipocyte fed on protonated D-glucose. This provides a reference for the characteristic peaks for TAG.

### Adipocytes

The time-dependence of the fat and glucose contents of the growth medium from cells undergoing adipogenesis and those of controls are shown in **Fig 4**. Control cells used for comparison here were grown in the same amount of glucose but without the aforementioned adipogenic reagents added. The total fat content remained essentially constant over more than a week after initiation of adipogenesis. The glucose level dropped over the same period. However, it remained at a sufficient level to maintain the cell culture (about 5.6 mmol/L). We can thus conclude that the cells always had sufficient material for lipid droplet formation from the medium itself, without having to resort to internal sources of precursors. It was important that extra D-glucose-d7 was not added to the medium, since the aim was to follow the spectral changes after a single event, the initiation of adipogenesis, without perturbing the system by subsequent changes in the medium.

**FIGURE 4.**
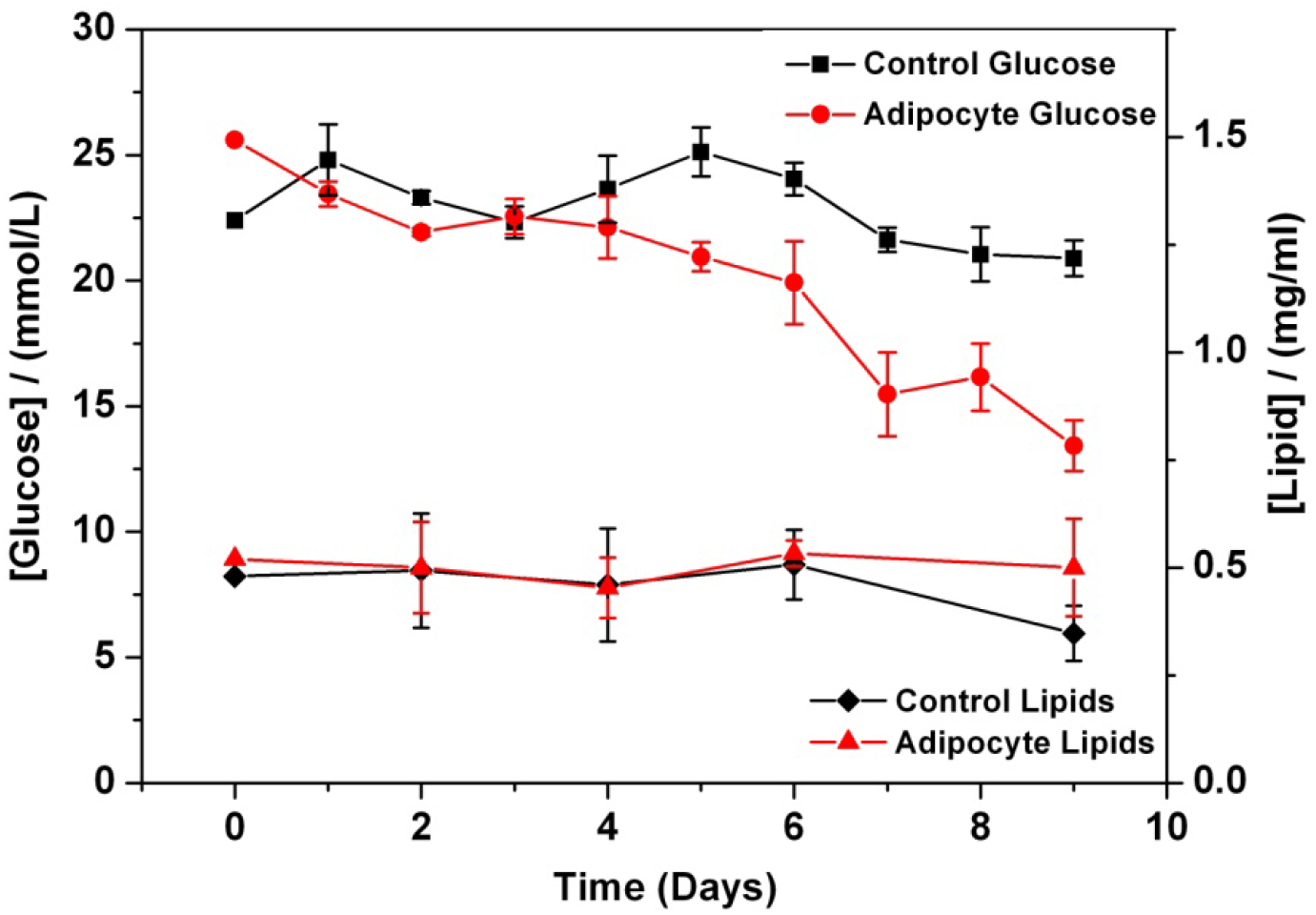
Glucose and fat content of medium during adipogenesis compared with control hMSCs (control glucose and control lipids) showing that the medium maintained sufficient levels of these precursors to sustain the formation of TAG.

Knowing that the cells had sufficient precursors for growth during the process of adipogenesis, the process itself could be studied in more detail over time. The most obvious features of the lipid droplet spectra that distinguish them from spectra of other parts of the cell are the bands at 1087 cm^−1^, 1306-1310 cm^−1^, 1751 cm^−1^ (carbonyl) and 2729 cm^−1^, as seen in **Fig. 3**. Furthermore, the lipid droplets display sharp bands at 2856 cm^−1^ and 2896 cm^−1^ which can be assigned to CH_2_ stretching; they can also be discerned as shoulders in the spectra of organelles, as organelles also contain lipid membranes that display similar CH_2_ stretching bands. As lipid droplet spectra are very easy to distinguish, because of the large contribution from CH_2_-stretching, mapping the positions of lipid droplets is relatively straightforward, as shown in **Fig. 5**.

**FIGURE 5:**
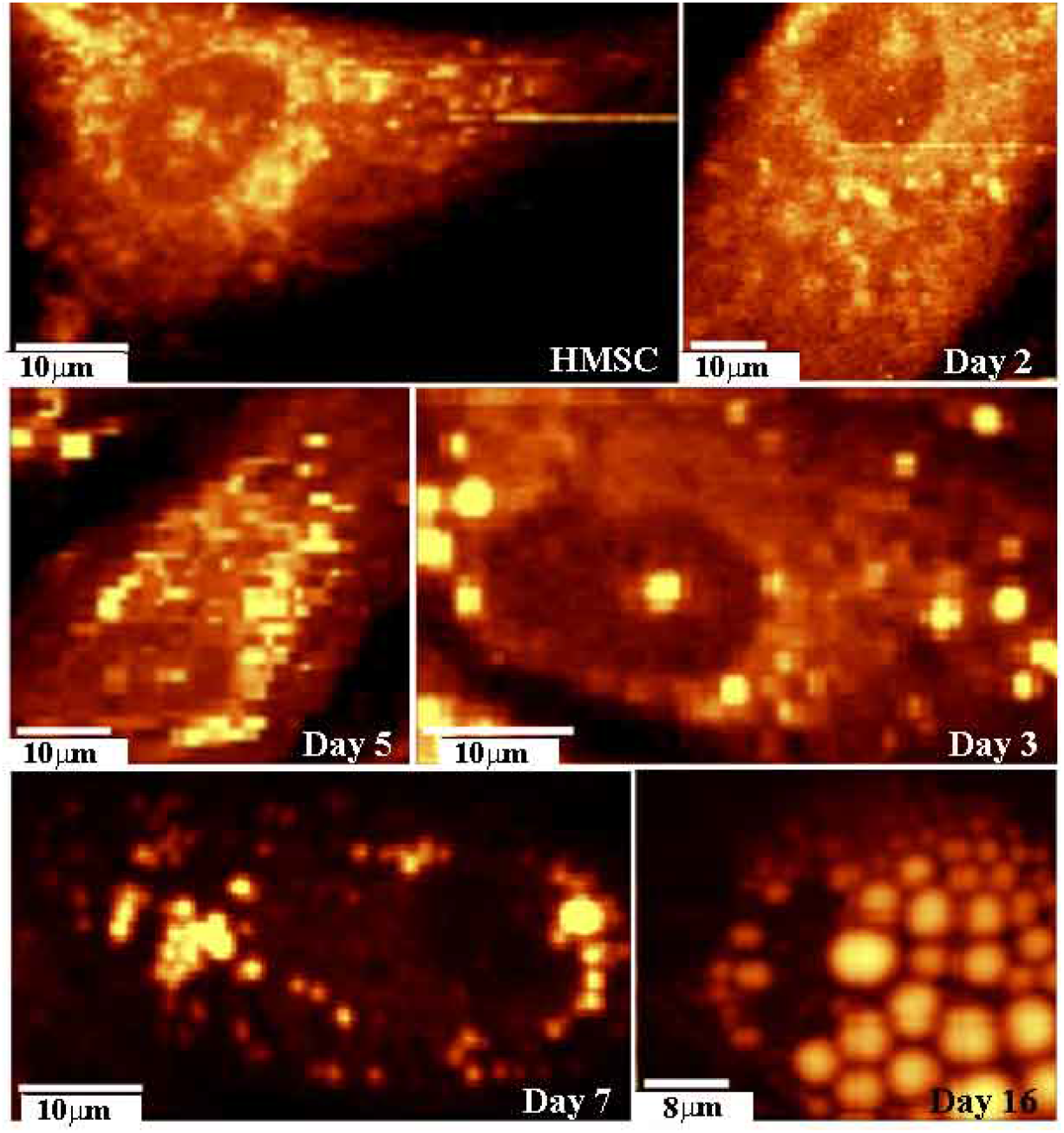
Typical Raman C-H stretch spectral maps of evolving adipocytes. Reproduced from (20) (permission granted from IEEE)

After differentiation was initiated, obvious lipid droplets became visible over several cells; they were readily apparent by day 3 and a small number of lipid droplets could be seen even by days 1 and 2, although the cell cultures were quite heterogeneous at all time points. Thus, poorly differentiated cells were apparent alongside well differentiated cells after 16 days in the same culture medium. The overall heterogeneity in the extent of adipogenesis day-by-day makes kinetic interpretations difficult. However, a general trend in the progress of the lipid droplet formation could be discerned in cells fed with non-deuterated glucose (**Fig 5**). However, it is apparent that at the earliest stages it was problematic to distinguish between lipids in organelles, vesicles and small lipid droplets. Thus, it was not possible to determine with certainty if a small lipid droplet was preformed prior to the initiation of adipogenesis, or whether it was formed after initiation.

Experiments were, therefore, conducted with D-glucose-d7 added to the culture medium in order to track the fate of glucose during its metabolism. Spectra of glucose d-7 in buffer and its metabolites taken from inside the adipocytes are shown in **Fig. 6**. D-glucose-d7 has C-D peaks at 2131 cm^−1^, 2169 cm^−1^ and 2232 cm^−1^, and has no bands in the C-H region. As soon as the glucose arrives inside the cell it is converted into its first metabolite, which can be seen in the cytoplasm as 3 peaks in the C-D stretch region, with maxima at 2128 cm^−1^, 2176 cm^−1^ and 2236 cm^−1^. This spectrum displays weak bands in the C-H region, probably due to protein co-located with the first metabolite in the cytoplasm within the probed volume. The most likely candidates for this intermediate are glucose 6-phoshate or fructose 6-phosphate, both of which can be expected to form quickly after the glucose enters the cell. Additionally fructose, glucose, (21) and glucose-6-phosphate (22) all have spectra comprising of 3 main peaks in the C-H stretch region that would shift to lower frequency by a factor of 1/√2 upon deuteration. Therefore, the observation of 3 peaks in the C-D region is consistent with this provisional assignment.

**FIGURE 6.**
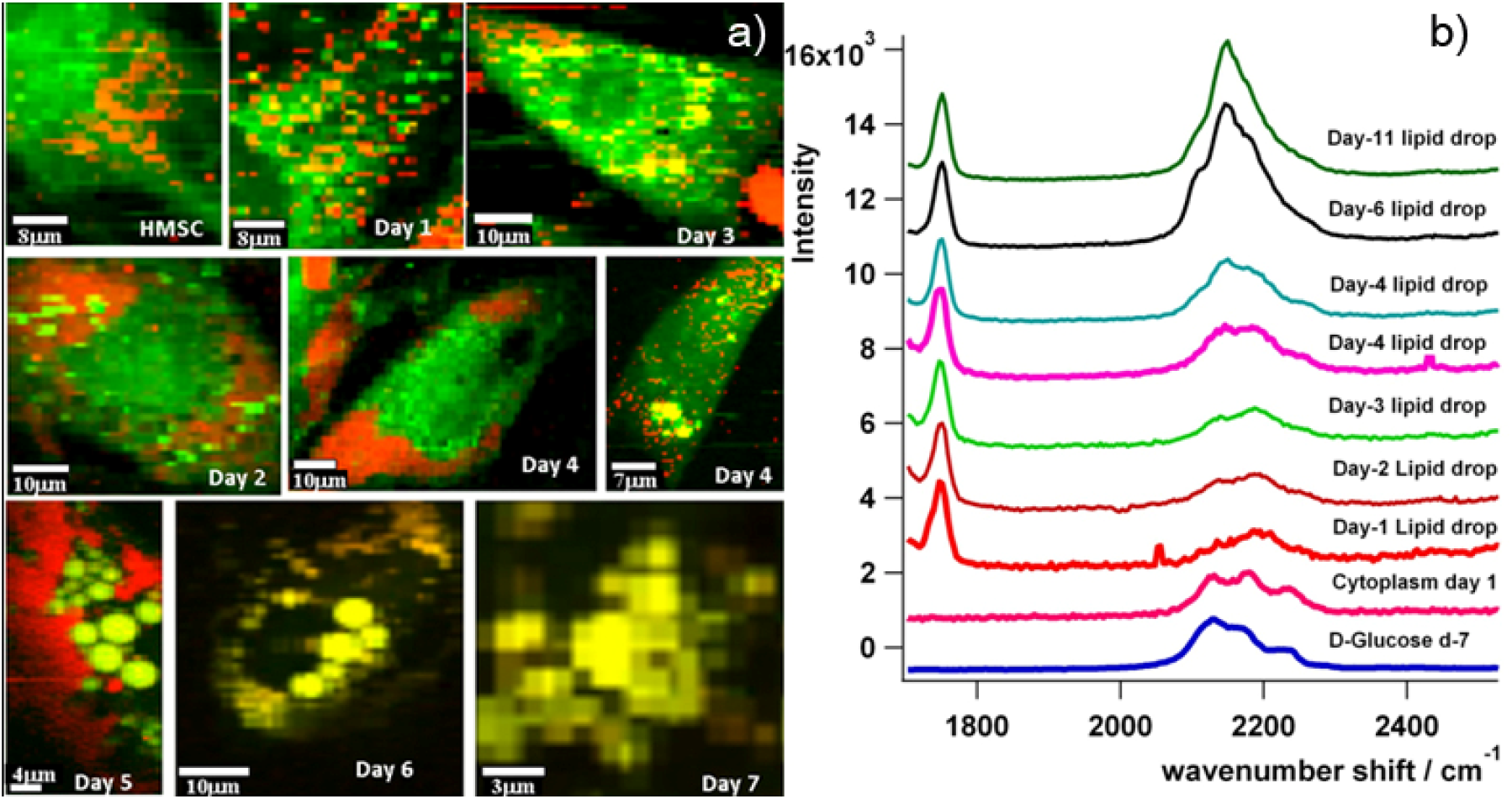
**a)** Time-course for adipogenesis and lipid drop formation reproduced from ref (20) (permission granted from IEEE); Red regions have strong C-D stretches Green regions have strong C-H stretches and yellow regions are lipid droplets with strong C-H and C-D stretches. **b)** Evolution of the C-D spectral region during adipogenesis.

As the adipogenesis proceeded, it became possible to identify small, induced putative lipid droplets, because they displayed spectra with significant peaks in both the C-H stretch region (2800 cm^−1^ to 3100 cm^−1^), due to fat chains, and in the C-D stretch region (2100 cm^−1^ to 2300 cm^−1^), due to the deuterated triacylglyceride backbone. Droplets that were preformed in the hMSCs before initiation of adipogenesis only displayed peaks in the C-H stretch region. By merging maps of the C-H and C-D stretch regions it was also possible to map the location of the different metabolites of the deuterated glucose in the cells over time (**Fig. 6**).

It was notable that the C-D stretch bands of the earliest lipid droplets from day 1 to day 3 were different to both the first metabolite in the cytoplasm, and to the spectra of the later C-D spectra in lipid droplets from days 6 to day 11. The earlier droplets were characterized by a central peak at 2187 cm^−1^ with shoulders at 2139 cm^−1^ and 2250 cm^−1^, whereas at later stages (e.g., day 6) during adipogenesis the spectrum of the C-D region is obviously modified and manifests as a sharp central peak at 2150 cm^−1^. This peak has 2 shoulders at 2109 cm^−1^ and 2172 cm^−1^.

In addition, the lipid droplet spectra display very strong C-H stretches and a prominent carbonyl peak, as do the non-deuterated lipid droplet spectra. Later still, by day 11, further slight modification occurred to the spectra, with the central peak still at 2150 cm^−1^, but with less prominent shoulders at 2105 cm^−1^ and 2239 cm^−1^.

From day 2 to day 6 it was also noted that there were often intermediate spectra with the later 2150 cm^−1^ peak pushed up through the spectrum definitive of earlier lipids. We could establish that these intermediate spectra could be nearly perfectly recreated and matched in terms of shape and intensity at all frequencies by mathematically mixing an early lipid spectrum from day 2 (designated here as “type 1”) with a later lipid spectrum from day 6 (designated as “type 2”). In contrast, the day 11 (designated as “type 3”), day 6 and day 1 spectra could not be created by mathematically mixing or stripping components of any of the other spectra, including the first metabolite. That led to the conclusion that there were 3 distinct classes of lipid droplet spectra in the C-D stretch region, designated types 1-3.

The spectra can be further analyzed, to confirm these initial conclusions, by fitting the C-D spectral region to Lorentzian curves. Glucose and its first metabolite clearly fitted well with a minimum of three Lorentzian peaks (**Fig 7 a** and **b**), as expected for glucose or fructose species. (21,22).

**FIGURE 7.**
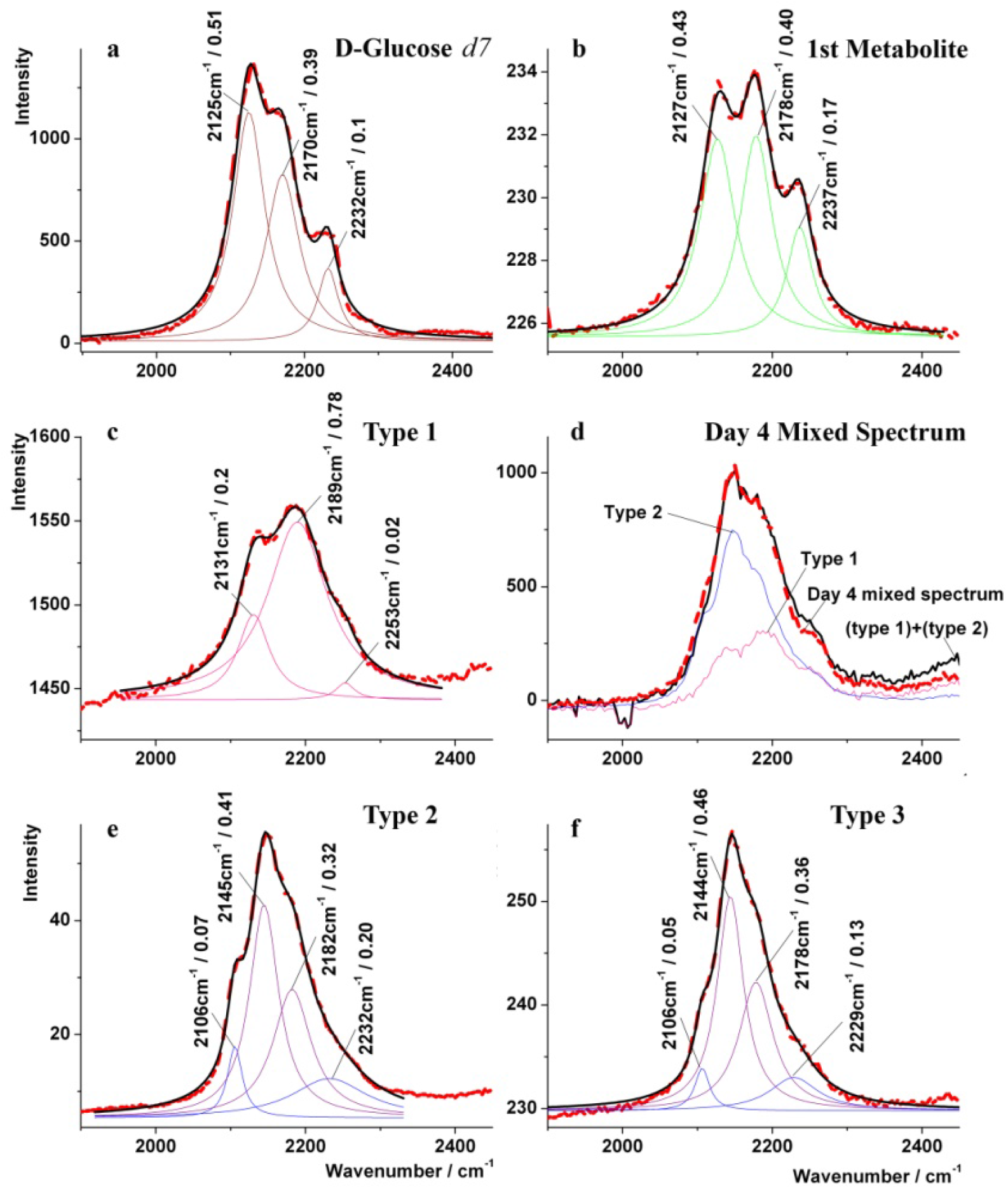
Lorentzian peak fitting to the C-D stretch region for glucose and its metabolites. The fits show that the initial C-D stretch peaks in the cytoplasm or not glucose, rather they are a glucose metabolite (a and b) type 1 spectra (c) from the early stages are clearly different to later spectra (e and f). Day 4 spectrum (d) is the result of a mixture of type 1 and type 2 spectra. Type 2 (e) and type 3 (f) spectra are most clearly differentiated by the differences in relative intensities of peaks, especially the peak at 2106 cm^−1^.

Type 1 lipid spectra (**Fig 7 c**) fitted to a minimum of three Lorentzian peaks, with one main peak at 2189 cm^−1^ dominating. Type 2 and type 3 lipid spectra (**Fig 7 e** and **f**) both fitted well to a minimum of 4 similarly positioned Lorentzian peaks. However, the ratio of these peaks was not the same for these two types, indicating that the peaks are due to different species, supporting the conclusions drawn from the initial analysis.

In particular, the peak forming the shoulder at 2106 cm^−1^ changes its relative contribution sufficiently to say that two species must be in coexistence under this spectral envelope. This supports the previous conclusion relating to the low frequency shoulder, which has its origin from the 2106 cm^−1^ peak. The day 4 spectrum (**Fig 7 d**) was not fitted, because the real situation would obviously require such a large number of Lorentzian peaks (7 peaks combined from type 1 and type 2 lipid drops) that the outcome would be unreliable. However, the comparison with the mixture of type 1 and type 2 spectra is shown, which reveals the origin of its spectral shape clearly to be from a mixture of the two types.

After such determination, it was possible to establish when each type of lipid spectrum occurs. To display this pictorially, a circle is divided into 3 parts, representing each type of spectrum, and then a dot is placed into each third of the circle, for each cell measured, whose lipid drop spectra most closely match the spectral type in that third of the circle. Where the spectrum is a mixture of types it is placed on the line between the defined regions. This is shown in **Fig 8(a)**. From this it can be seen that the lipid spectra start as type-1 spectra and gradually evolve through type-2 to type-3 spectra.

**FIGURE 8.**
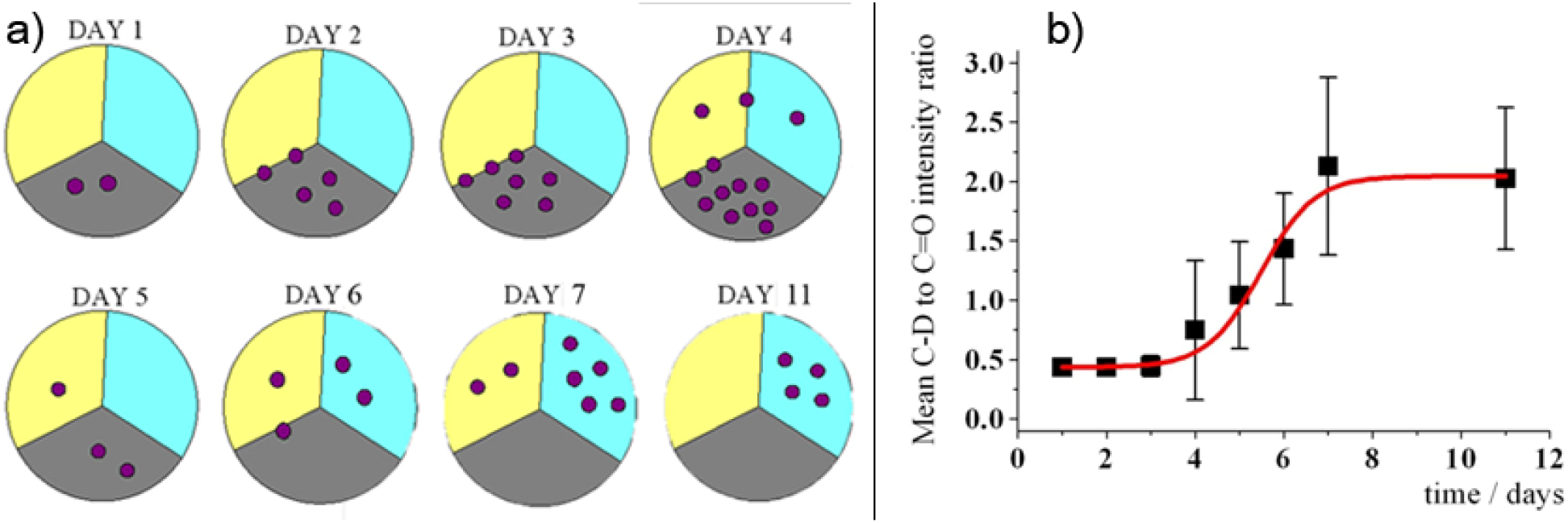
**a)** Spectral evolution plotted as spectral type (grey region = type 1, Yellow region = type 2 and blue region = type 3 spectra). Each spot represents an instance where that type of spectrum was found in the lipid droplets in a cell. Each spot represents analysis of a different cell’s Raman map. **b)** The ratio of carbonyl to C-D stretch intensity plotted against time. Data is pooled from several images of cells (as represented by the number of spots in a))

Also plotted in **Fig. 8(b)** is the evolution of peak intensity ratios for the carbonyl stretch region and the C-D stretch region. The carbonyl intensity can be taken as a reliable reference point because it is a constant for every fatty acid chain that adds to the TAG-backbone. It can be seen clearly from both the spectra in **Fig. 6** & **7** and the time-course in **Fig. 8** that a spectral evolution in terms of band intensities occurs together with the changes in the spectral shape and type.

Although a full assignment of the source of this spectral evolution is beyond the scope of this study, it is possible to surmise conditions that could lead to the observations made here. Thus, there are important implications from the observation of evolution in Raman spectra in lipid droplets in the C-D stretching region that at least show that lipid composition is not constant over time in terms of the deuteration pattern of the glucose fragments forming the TAG backbone in the droplets. However, this variation can have at least two possible sources, in view of the overall complexity of the TAG synthesis pathways.

A first possibility is that lipids are undergoing some chemical modification inside the droplets over time, eventually leading to TAG, and that there are transient chemical forms that result in the observed spectral evolution. An example is diacylglycerol transacylase, responsible for 1,2-diacyl-sn-glycerol synthesis, which might also catalyse further acylation reactions to form triacyl-sn-glycerol (23,24). Although diacylglycerol transacylase is an ER membrane protein, the exact location of where this process occurs is not known, but may to some extent be in or around lipid droplets. This is particularly relevant, if the putative lipid droplets do in fact bud off from the ER in the first place, with the ER membrane forming the membrane of the droplet (5). It is also possible that within the lipid droplets, post-processing of TAGs may occur.

The second possibility is that there are different pathways for deuterated TAGs arrival into the lipid droplet, resulting from different metabolic pathways for d7-D-glucose. For example, in adipocytes, two pathways from glucose to TAG are already known.

In the Kennedy pathway (also known as the sn-glycerol-3-phosphate pathway), (25) glucose is converted to glucose-6-phosphate, fructose-6-phosphate, glycerol-3-phosphate, lysophosphatidic acid, diacylglyceride, and then TAG. This pathway leads to various deuterium substitution sites on the TAG backbone (20). The main source of the glycerol-3-phosphate in this pathway is the catabolism of glucose, although some can come indirectly from glyceroneogenesis via pyruvate, and this may be the main source in adipose tissue (26).

Pyruvate can itself come from the same deuterated glucose source as the glycerol-3-phosphate, but one would not necessarily expect the same deuteration pattern on the TAG backbone. Unfortunately, to absolutely assign the spectral changes observed to definite chemical structures and pathways is again beyond the scope of the current work, and would involve synthesis of some model deuterated compounds and measurement of their spectra for comparison with the spectra considered here.

One additional point of interest is that the d7-D-glucose metabolites were only positively observed in the cytoplasm and in the lipid droplets themselves. There was no observed accumulation of metabolites in any other part of the cell, nor was there localization in any other form having a different spectrum from the 4 metabolites identified. This can be taken to mean that either the other sites for this process do not exist, or at least not in the cells examined here, or that they do exist, but the turnover of the metabolites is so fast that they do not accumulate in sufficient concentration to be observed in that location. In this regard, images such as the Fig. 6 day 5 Raman map are intriguing, as they show the cytoplasm awash with the D-glucose-d7 first metabolite surrounding a cluster of lipid droplets containing a mixture of type 1 and type 2 lipid. However the glucose metabolite appears to be depleted closest to the lipid droplets. This depletion may imply that the first metabolite is being consumed within the lipid droplets themselves, or on their surfaces.

Cluster analysis of Raman maps for adipocytes are shown in **Fig. 9**. Spectra derived from cluster analysis confirmed the spectral shapes shown in **Fig. 6** for both early putative lipid drops and for the later lipid drops. It also matches well with the merged C-D and C-H stretch maps in **Fig 6**. Additionally the spectra of lipid droplets made with deuterated glucose are very similar to the spectra of droplets made with protonated glucose, except in the C-D stretch and C-H stretch regions. This is shown in **Fig. 10**.

**FIGURE 9:**
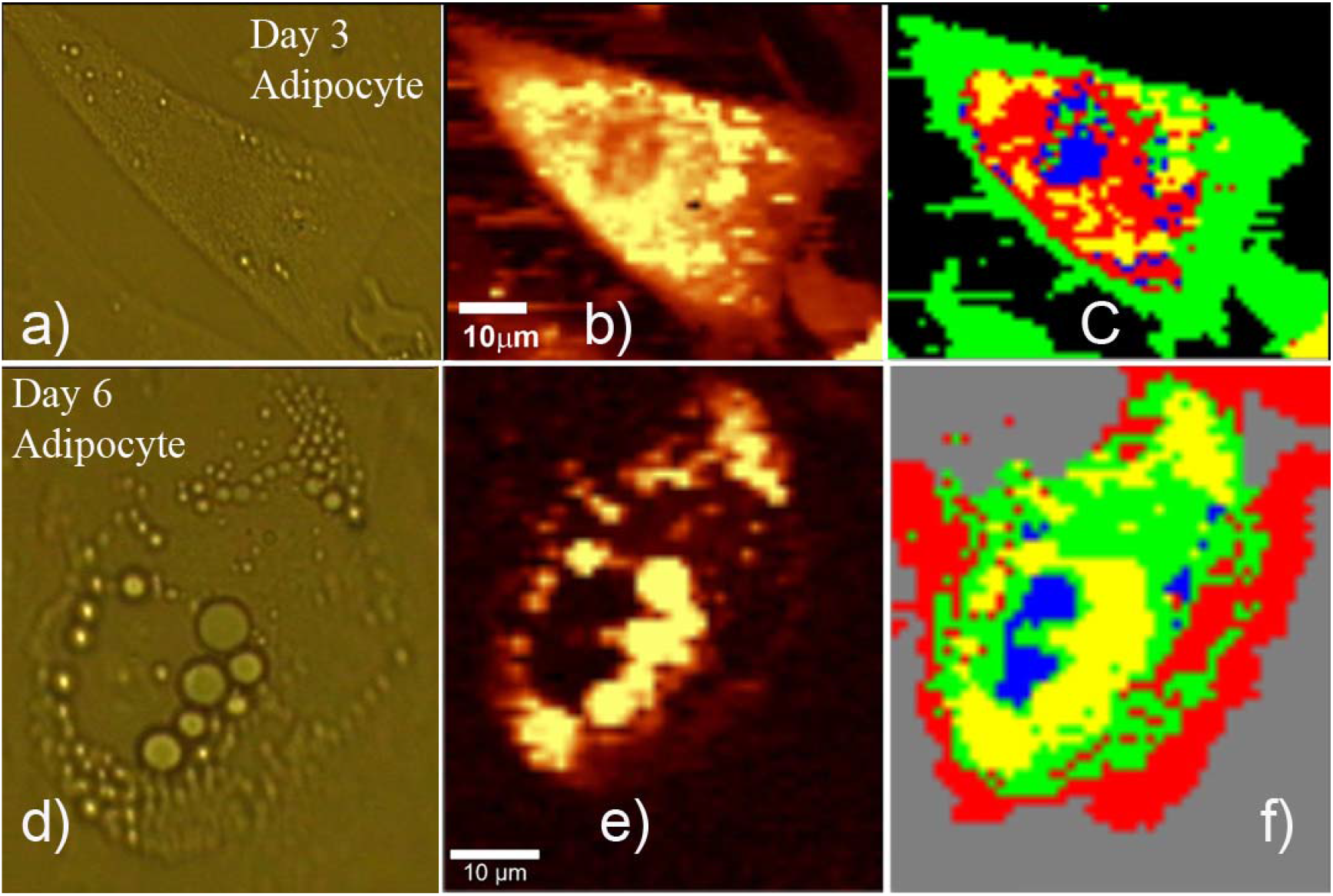
Top **a)** Optical image, **b)** Raman C-H stretch map and **c)** Cluster analysis of a 3-day old adipocyte fed on deuterated glucose. Yellow is the first glucose metabolite, and early stage lipid droplets, respectively derived from cluster analysis in the C-D region. Green represents less glucose rich areas, with the nucleus (blue) derived from a spectral cluster analysis from 500cm^−1^ to 3100cm^−1^. Bottom **d)** Optical image, **e)** Raman C-H stretch map and **f)** Cluster analysis of the Raman map based on the C-D stretch region (yellow in (e)) with (in the cluster map) blue showing the nucleus, yellow again showing lipid droplets, green as cytoplasm displaying significant C-D from lipids and red showing areas of cytoplasm showing the least C-D bands. Similar cluster maps could be derived from analysis between from 500cm^−1^ to 3100cm^−1^.

**FIGURE 10.**
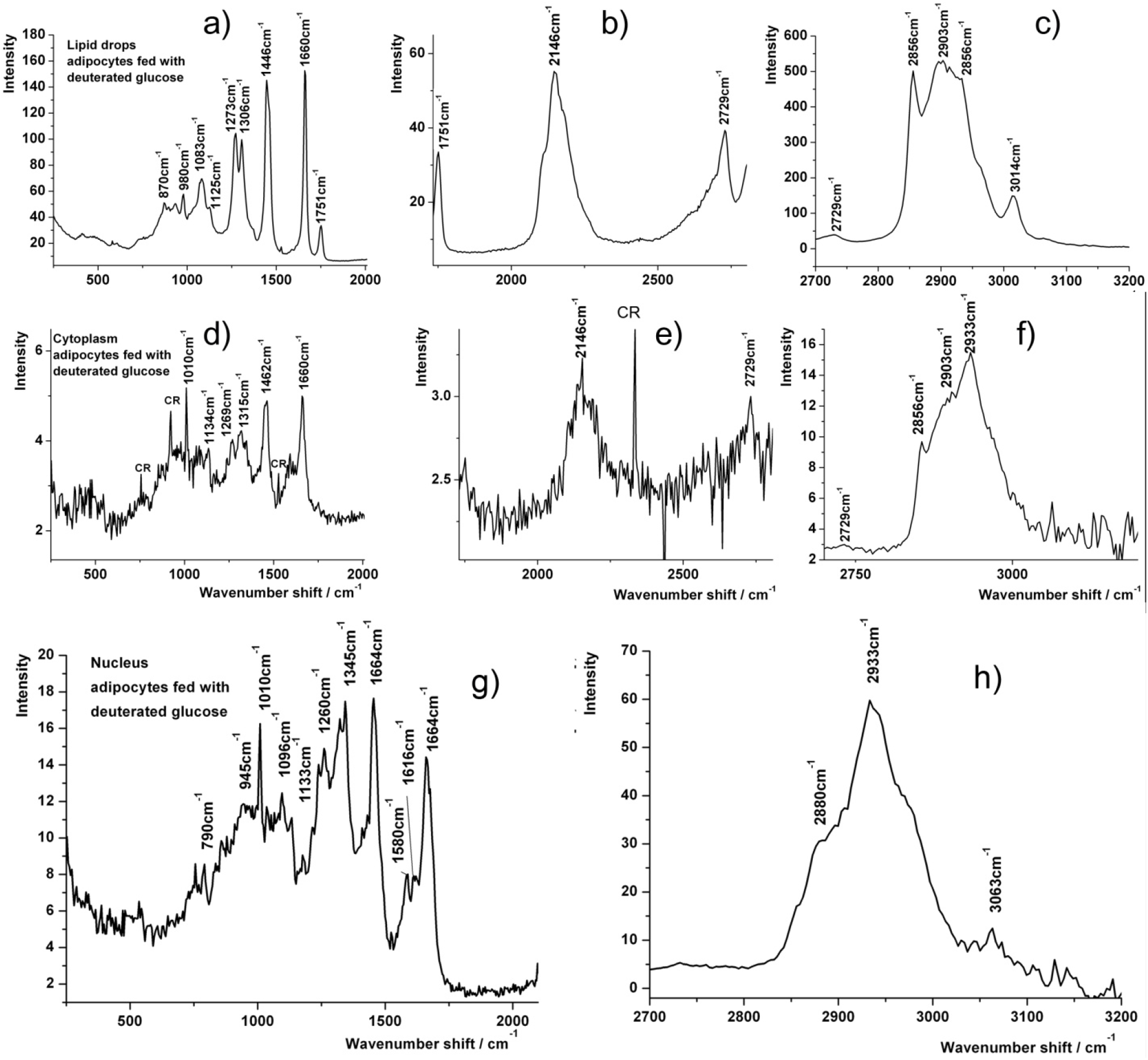
Top panel: **a) – c)** Spectra of yellow clusters (lipid droplets) from the cluster analysis of the day 6 adipocyte in Fig. 9(f). Middle panel: **d) – f)** Spectra derived from cluster analysis of day 6 adipocyte (figure 9(f)) from regions far away from any visible lipid droplets. CR=cosmic ray. The appearance of clear deuterated TAG peaks far from the lipid droplets demonstrates the presence of significant levels of TAG in the cytoplasm. Bottom panel: **g)** and **h)** Spectra from the nucleus of a day-6 adipocyte.

A notable feature of the day 6 spectra (Fig. 6) was that lipids appeared to be ubiquitous about the cell. Apart from within the nucleus, it was not possible to find a cellular region, even in the parts of the cytoplasm farthest away from any visible lipid droplets, which did not show a characteristic lipid C-D stretching peak.

This implies that lipids are being transported throughout the cytoplasm, either at their own solubility limit or mediated by lipid-binding proteins or vesicles that are too small to see individually by optical means. Additionally, the C-D spectra observed in the cytoplasm were essentially the same as those in the visible droplets. This finding may have implications for the mechanism of lipid droplet growth.

It has always been known that droplet fusion is a rather rare event (5). However, lipid droplets are known to grow larger over time. It is, therefore, possible that the droplet coarsening mechanism in adipocytes may be related to the evaporation condensation models of droplet coarsening that are found in other more physical systems. In the latter case, small droplets of high curvature evaporate and transport through the cytoplasm to condense into larger droplets of lower curvature, in this case possibly mediated by a transport agent such as a lipid-binding protein (27). Alternatively, this may show transportation of TAG from a region where it is generated to the lipid droplets, although there is no identifiable point of origin, as discussed above.

Another significant point in relation to the evolution of Raman spectra during adipogenesis is that there are peak ratio changes comparing the carbonyl stretch (which is constant in proportion for every fat chain added) and the 1660 cm^−1^ C=C stretch. Additional band ratio changes were seen for 1265 cm^−1^ and 1306 cm^−1^ which are for =C-H and −C-H stretches, respectively (28). It was observed that the bands resulting from unsaturated fats were represented in higher proportion in the earlier spectra of lipid drops than they were at later times. The trend can be seen in **Fig. 11** and with reference to **Table 2**.

**Table 2.** Band assignments for lipid droplet spectra.

|  |  |  |  |  |  |
| --- | --- | --- | --- | --- | --- |
| 1086 | C-C | 1265 | =C-H | 1306 | -C-H |
| 1448 | -C-H | 1655 | C=C | 1753 | C=O |
| 2855 | -C-H | 2894 | -C-H | 2925 | -C-H |

**FIGURE 11.**
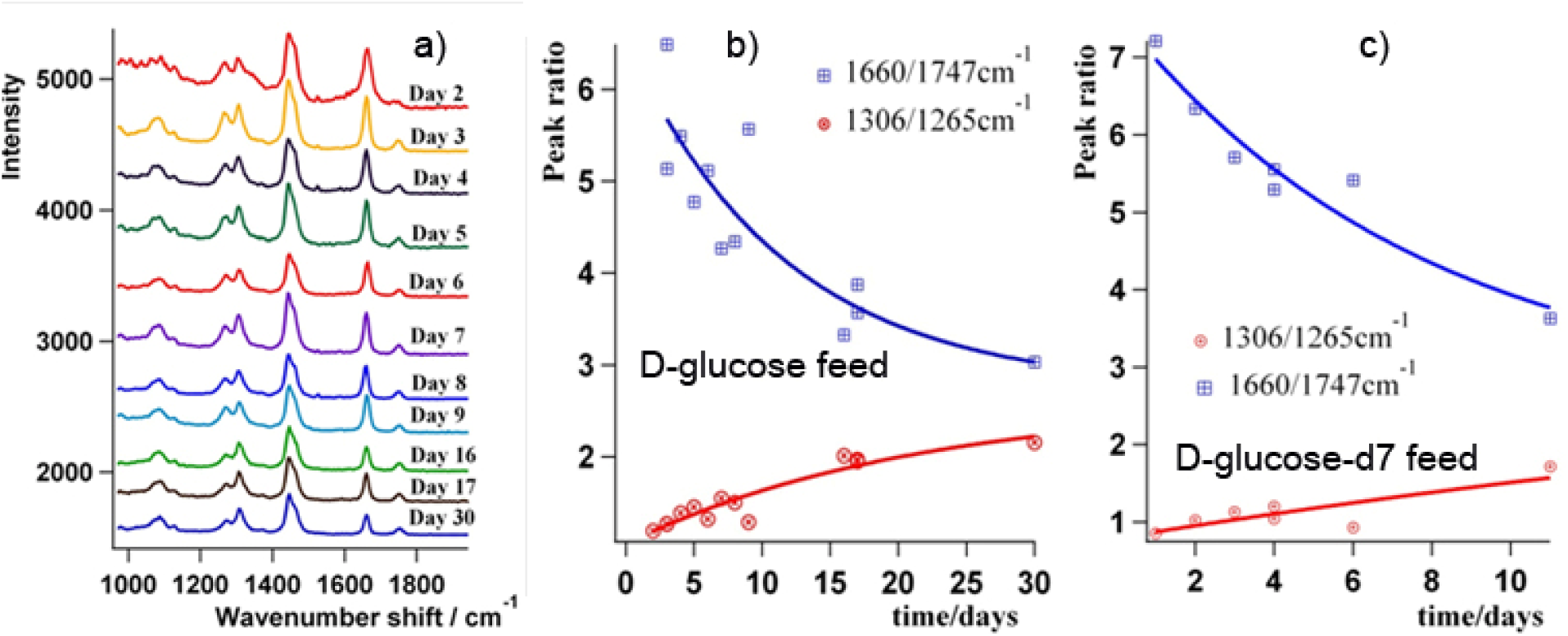
Right: **a)** Spectral evolution of Raman bands over time (adipocytes fed on D-glucose). Left: Ratios of peaks corresponding to saturated and unsaturated fats over time for adipocytes fed on **b)** D-glucose and **c)** D-glucose-d7. Relative spectral changes at 1306 cm^−1^, 1265 cm^−1^, 1660 cm^−1^ and 1747 cm^−1^ indicate the faster arrival of unsaturated fats in the lipid droplets compared to saturated ones. All data are from pooled measurements of several cells on different days as for Fig 8 a) and b).

This implies that, overall, unsaturated fats arrive in the lipid drops more efficiently than the saturated fats. It is interesting to note that adipocyte fatty acid-binding proteins have been reported to have higher affinity for unsaturated fats (oleic acid) than for saturated ones (stearic acid) (29, 30). However, it must be noted that the overall arrival time of the different states of fat saturation will also depend on other parameters, such as their intrinsic solubility.

## III. CONCLUSIONS

Adipogenesis was initiated in hMSCs using D-glucose and D-glucose-d7. Four distinct metabolites were identified leading to the final triacylglyceride. These metabolites were mapped inside the cells over time. The first metabolite is presumably glucose-6P or fructose-6P. The second and third metabolites could be intermediate species on their way to the final metabolite, which is assigned as TAG; for example, the conversion of diacylglyceride to TAG, or the post-synthesis modification of TAG. Alternatively, it might indicate that there are different metabolic pathways leading from D-glucose-d7 to TAG. It is certainly conceivable that the evolution in spectra is caused by a combination of the above.

It can be posited that the TAG backbones in the droplets have a mixture of forms ranging from 2-4 deuterium atoms on the TAG backbone and that the pattern of deuteration would depend on the pathway. Another important finding is that the proportion of unsaturated fats in lipid droplets is higher at the beginning of the process of adipogenesis than it is at the end.

This may be due to the relative affinity of saturated and unsaturated fats for lipid-binding proteins. Finally, the levels of TAG found in the cytoplasm of adipocytes, at intermediate times (day 6), indicates that TAG is mobile in the cytoplasm at measurable levels, suggesting that an evaporation condensation model for droplet coarsening is possible, as TAG can be expected to condense into larger droplets in preference to smaller ones. The work here shows the value of Raman imaging as a means of understanding biochemistry in the living cell.

## ACKNOWLEDGMENTS

This work was funded by A*STAR JCO. J. H. and G. S. S are grateful to Shawn Lee, Witec Pte Ltd for the use the Raman system from Witec (Ulm Germany). DGF was funded by the North West Cancer Research Fund. SG was funded by ASTAR-JST.

## FOOTNOTES

The abbreviations used are:

TAG: triacylglyceride
CARS: Coherent Anti-Stokes Raman scattering
hMSCs: human Mesenchymal Stem Cells
D-glucose-d7: D-Glucose-1,2,3,4,5,6,6-d7, 97 atom % D
ER: Endoplasmic Reticulum
glucose-6P: glucose-6-phosphate
fructose-6P: fructose-6-phosphate.

